# The power of model-to-crop translation illustrated by reducing seed loss from pod shatter in oilseed rape

**DOI:** 10.1101/604769

**Authors:** Pauline Stephenson, Nicola Stacey, Marie Brüser, Nick Pullen, Muhammad Ilyas, Carmel O’Neill, Rachel Wells, Lars Østergaard

## Abstract

In the 1980s, plant scientists descended on a small weed *Arabidopsis thaliana* (thale cress) and developed it into a powerful model system to study plant biology. The massive advances in genetics and genomics since then has allowed us to obtain incredibly detailed knowledge on specific biological processes of Arabidopsis growth and development, its genome sequence and the function of many of the individual genes. This wealth of information provides immense potential for translation into crops to improve their performance and address issues of global importance such as food security. Here we describe how fundamental insight into the genetic mechanism by which seed dispersal occurs in members of the Brassicaceae family can be exploited to reduce seed loss in oilseed rape (*Brassica napus*). We demonstrate that by exploiting data on gene function in model species, it is possible to adjust the pod-opening process in oilseed rape thereby significantly increasing yield. Specifically, we identified mutations in multiple paralogues of the *INDEHISCENT* and *GA4* genes in *B. napus* and have overcome genetic redundancy by combining mutant alleles. Finally, we present novel software for the analysis of pod shatter data that is applicable to any crop for which seed dispersal is a serious problem. These findings highlight the tremendous potential of fundamental research in guiding strategies for crop improvement.

**Keymessage:** Elucidation of key regulators in *Arabidopsis* fruit patterning has facilitated knowledge-translation into crop species to address yield loss caused by premature seed dispersal (pod shatter).

## Introduction

Food security is being challenged worldwide due to population increase and climate change (Tilman et al. 2011; Fischer et al. 2014; Hunter et al. 2017). It is therefore crucial that plant scientists explore every opportunity for enhancing the efficiency with which crops are grown. Advances in plant molecular biology, genetics and genomics have over the last three decades led to an explosion in our understanding of specific biological processes, the function of individual genes and genome dynamics during plant development. This wealth of information holds an incredible potential to be exploited for translation into improved and sustainable crop production (Boden, Østergaard 2019).

More research funds have been invested into understanding the biology of *Arabidopsis* than to any other plant species in the belief that many processes controlling all aspects of plant development are conserved across the plant kingdom. Attempts to transfer knowledge from *Arabidopsis* to crops seems a valid test of this assumption. Since members of the *Brassica* genus are the closest crop relatives to *Arabidopsis*, model-to-crop translation between the *Arabidopsis* and *Brassica* crops provide potentially huge opportunities. The aim of this paper is to provide an illustrative example of how understanding the molecular details of a specific process during plant development can be used to address a serious barrier to more efficient crop production.

### Morphological conservation of fruit structures in *Arabidopsis* and *Brassica* species

*B. napus* along with other important *Brassica* crops is a member of the Brassicaceae family, which also contains *Arabidopsis*. The close evolutionary relationship between them is reflected in the conservation of the genome structure with segments of genetically linked loci of the *B. napus* genome corresponding to regions of the *Arabidopsis* genome exhibiting a high level of synteny (Parkin et al. 2005). Moreover, *B. napus* and other *Brassica* species have very similar plant architecture and organ morphology to *Arabidopsis* (Fig. 1) and it is therefore plausible that knowledge of significant biological processes obtained in *Arabidopsis* could be used to improve the performance of oilseed rape.

**Fig. 1.**
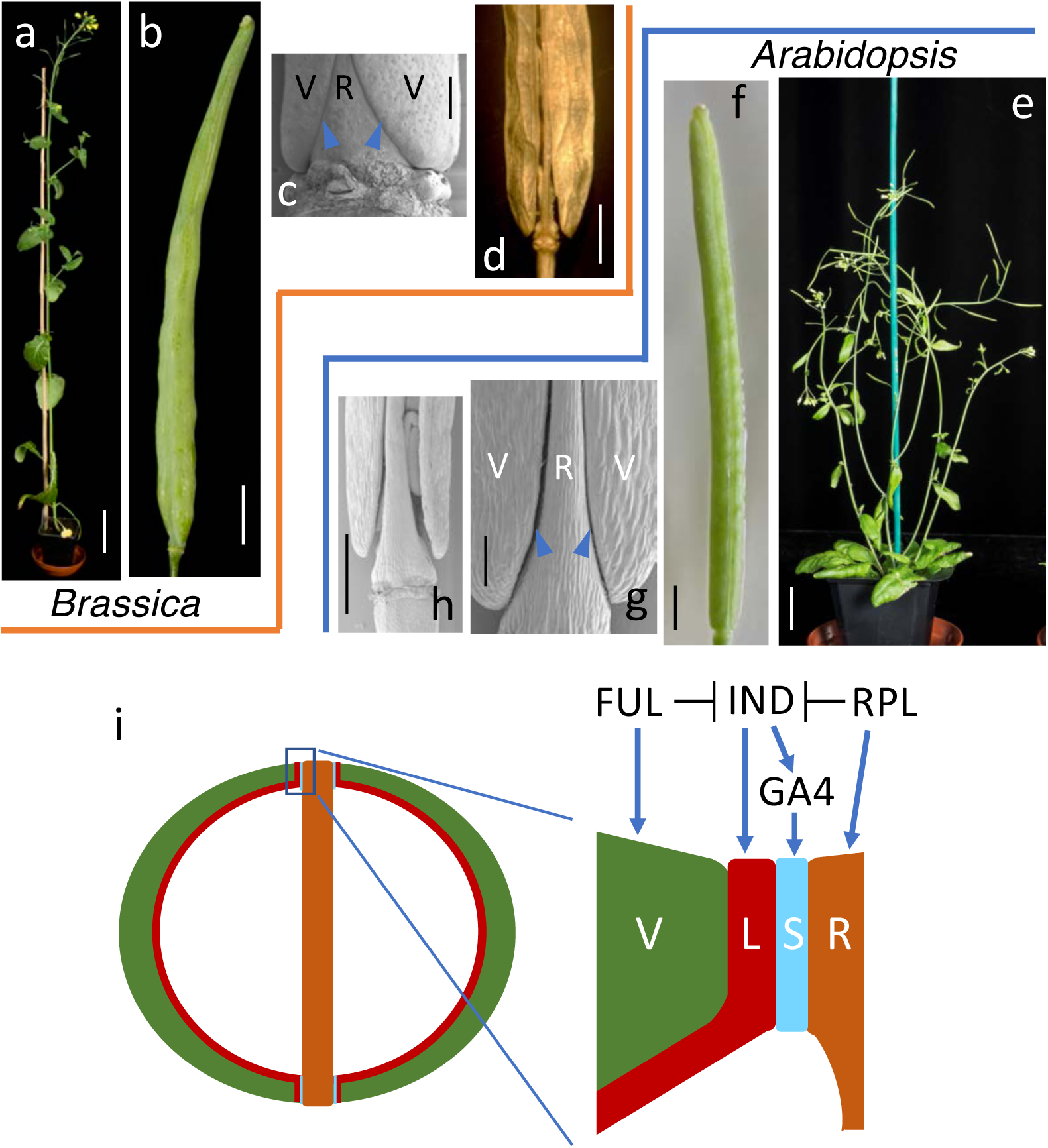
Conserved plant architecture and pod morphology between *Brassica* (**a-d**) and *Arabidopsis* (e-h). **a** 8-weeks old *Brassica rapa* plant. **b** mature pod (stage 17) from *B. rapa*. **c** SEM of stage-15 *B. rapa* pod. **d** wholemount image of dehiscing *B. napus* pod (stage 19). **e** 5-weeks old Arabidopsis plant. **f** mature pod (stage 17) from Arabidopsis. **g** SEM of stage-17 Arabidopsis pod. **h** SEM of dehiscing *Arabidopsis* fruit (stage 19). **i** Schematic cross section of pod from Arabidopsis/Brassica with valves in green, replum in brown, lignified layer in red and separation layer in light blue. A simplified regulatory network is indicated above the close-up on valve margin region. Blue arrowheads in **c** and **g** indicate position of valve margins. V valve, R replum, L lignified layer, S separation layer. *Scale bars* correspond to 100 µm in **g**, 500 µm in **c** and **h**, 1 mm in **f**, 5 mm in **d**, 1 cm in **b**, 2 cm in **e** and 10 cm in **a**.

Pod shatter is a term used for the preharvest fruit opening and seed dispersal of oilseed rape (*B. napus*) leading to yield loss. The similarities in fruit morphology between *Arabidopsis* and *Brassica* species (Fig. 1) suggest that the underlying patterning mechanism is conserved. It is therefore possible that knowledge on fruit opening in *Arabidopsis* can be exploited to address the problem of pod shatter in oilseed rape. After fertilisation at developmental stage 13 (stages defined for *Arabidopsis* flower development in Smyth et al. 1990 and conserved in Brassicas as described in Girin et al. 2010), Brassicaceae fruits elongate and form a number of specialised tissues including valves (also known as pod walls), a central replum and valve margins that form at the valve/replum borders where fruit opening will take place (Fig. 1c and 1g). This process is the result of precise tissue-and cell-type specification allowing the separation of the valves from the replum and leading to the timely dispersal of mature seeds (Fig. 1d and 1h; Roberts et al. 2002). The valve margins are composed of two distinct cell types, a lignified layer and a separation layer (Fig. 1i). Upon fruit maturation, cells in the valve margins mediate fruit opening through degradation of the pectin-rich separation layer by secreting polygalacturonase enzymes (Ogawa M et al. 2009; Petersen et al. 1996; Degan et al. 2001; Spence et al. 1996).

Although efficient seed dispersal is an advantage for plants growing in the wild, unsynchronised pod shatter of oilseed rape causes average annual losses above 10% of harvest (Price et al. 1996) and can exceed 70% under adverse weather conditions. Controlling pod shatter is in fact such an important trait for oilseed rape yield increase, that many other traits focused around plant architecture are aimed at reducing pod shatter-mediated seed loss in the field (Morgan et al 2000).

### Conservation of genetic and hormonal activities in Brassicaceae fruit development

Setting up the overall patterning process of the *Brassicaceae* fruit is a prerequisite for proper development. Several of the key genetic factors from *Arabidopsis* have been identified and the interactions between them established (Fig 1i). For a comprehensive description of all known components, see Dinneny et al (2005). Here, we focus on the *INDEHISCENT* (*IND*) gene, which encodes a basic helix-loop-helix (bHLH) transcription factor that is essential for valve margin formation (Liljegren et al. 2004). The highly specific expression of the *IND* gene in valve margin tissue is ensured by the repressing activities of the *FRUITFULL* (*FUL*) gene in the valves and the *REPLUMLESS* (*RPL*) gene in the replum (Ferrándiz et al. 2000; Roeder et al. 2003; Dinneny et al. 2005).

We previously found that this genetic network interacts with activities of the phytohormones auxin and gibberellin (GA) to ensure proper fruit patterning in *Arabidopsis* (Sorefan et al 2009; Arnaud et al. 2010; Girin et al. 2011). In wild-type fruits, both local depletion of auxin and biosynthesis of GA are required at the valve margin for specification of the separation layer where fruit opening takes place. IND mediates these events by directly regulating genes involved in both processes. The *GA4* gene in Arabidopsis encodes an enzyme (GA3OX1) that mediates the final step in the biosynthesis of active gibberellins (GA_1_ and GA_4_) (Talon et al. 1990), and our previous work revealed that IND binds to and directly induces expression of the *GA4* gene (Arnaud et al. 2010).

Yield of oilseed rape could be significantly improved by controlling pod shatter. Studies of fruit development in *Arabidopsis* and diploid *Brassica* species have provided directions for achieving this. For example, we have shown that ectopic expression of the *Arabidopsis FUL* gene under the *CaMV 35S* promoter leads to a complete loss of shattering in *B. juncea* as previously observed in Arabidopsis (Fig. 2; Østergaard et al., 2006; Ferrándiz et al. 2000). Although this result demonstrates that it is possible to transfer knowledge from the model system to *Brassica* crops, total encapsulation of the seeds in a fruit with no remaining ability to dehisce is undesirable for oilseed rape production, as significant losses would be incurred when attempting to retrieve the seeds. Subsequently, we used *B. rapa* (a progenitor of *B. napus*) as a system to directly target an *IND* homologue (*BraA.IND.a*), which exists as a single-copy gene in the *B. rapa* genome. Using a *B. rapa* TILLING population (Stephenson et al. 2010), we demonstrated that strong *braA.ind* mutant alleles are indehiscent (Fig. 2b) and that it was possible to obtain weaker alleles in which valve margin tissue was only partially lost (Girin et al. 2010). This intriguing result showed that it is feasible to fine-tune the level of shatter resistance in *Brassica* fruits by adjusting the activity of *IND* homologues. We also identified a functional *IND* homologue, *BolC.IND* in the other *B. napus* progenitor, *B. oleracea* (Girin et al. 2010). Moreover, in agreement with *GA4* being positively regulated by *IND* as described above, we also demonstrated that a gene-edited line in which two *BolC.GA4* paralogues were knocked out resulted in indehiscent (i.e. pod shatter resistant) pods (Fig. 2b; Lawrenson et al. 2015).

**Fig. 2.**
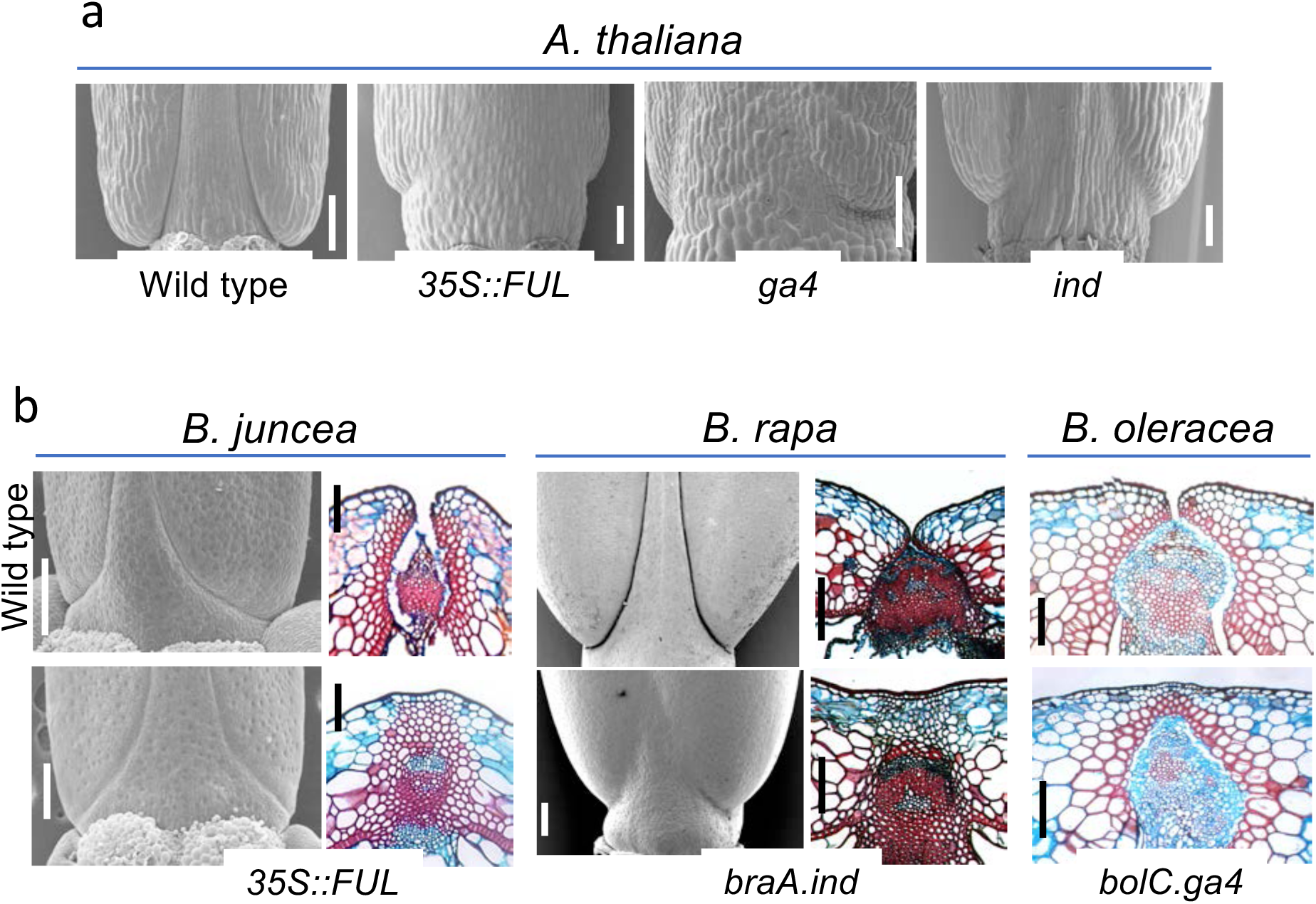
Effect of gain-and loss-of function of genes involved in valve margin formation. **a** SEM images of the bases of fruits at stage 17 from *Arabidopsis* wild type, *35S::FUL, ga4* and *ind* mutants. **b** SEM images and staining of cross sections are shown for the indicated *Brassica* species. Sections are from the middle of the fruits where the replum is narrower. Upper row display images from wild type fruits at stage 17 whereas stage-17 fruits from the indicated mutant genotypes are shown in the lower row. *Scale bars* correspond to 100 µm for *A. thaliana* SEMs and *B. juncea* and *B. oleracea* sections, 200 µm for *B. rapa* sections and 250 µm for *B. juncea* and *B. rapa* SEMs.

In this study, we obtain and analyse mutant combinations of *IND* and *GA4* paralogous genes of the allotetrapoid *B. napus* (oilseed rape). Our results provide the final stage of a model-to-crop translation pipeline from fundamental discoveries in a model system via proof-of-concept in diploid *Brassica* species to validation in oilseed rape thereby revealing the power of such strategies for crop improvement.

## Materials and Methods

### Plant material and plant growth

*B. napus* var. Cabriolet wild type and mutants were sown in F9 pots containing F1 compost and then grown at long day (16 hrs light/8 hours dark) at 18°C day and 12°C night temperature. After 6 weeks vernalisation at 5°C, plants were potted on into 1-L pots containing John Innes Number 2 compost. They were then grown in a glasshouse with long day conditions at 18°C day and 12°C night temperature.

### Phylogenetic analyses

*Brassica* IND and GA4 sequences were identified from EnsemblPlants by using BLAST with the *Arabidopsis* IND and GA4 protein sequences, respectively. Alignments were produced using the ClustalOmega software from EMBL-EBI and phylogenies were produced using the phylogeny.fr software package (Dereeper et al. 2008).

### Random Impact Test (RIT) assay

Pod shatter resistance was quantified by the RIT assay first described in Morgan et al. (1998). Fruits were harvested at stage 19 (stages defined in Smyth et al. 1990 and Girin et al. 2010), when they were fully dry, but unopened and placed in portions of twenty into small perforated bags, which allow moisture exchange. Bags of fruits were placed in a dehumidification chamber and equilibrated to 50% humidity for >48 hours. Twenty undamaged fruits were used in each separate assay and assays were performed in triplicates (*i.e.* 60 fruits required per data point). Fruits were placed together with six 8-mm steel balls in a 20-cm-diameter cylindrical container. The container was shaken in the Random Impact Test machine at a frequency of 4.98 Hz and a stroke length of 51 mm for 8-sec intervals until all the fruits were dehisced. After each interval, fruits were examined and the number of intact fruits remaining were counted. Fruits that had released any of their seeds were considered broken.

### podshatteR software

Calculation of RIT_50_ values were performed using the podshatteR software developed in this study. A description and step-by-step guide is provided in the Online Resource material. Briefly, this software provides an upgrade in curve fitting compared to previous reports (Morgan et al. 1998). Following the initial plotting of data by podshatteR, data was visually inspected to identify any errors or ambiguities. Curve fitting was then performed using an exponential decay function to keep the number of free parameters minimal i.e.

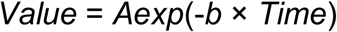

where A is the intercept (which should be similar to the starting number of pods), and b is the decay rate. Through propagation of errors confidence intervals were computed for each sample. The half-life was then calculated for each sample as the value at which half the initial number of pods has shattered, following the model fit.

## Results and Discussion

### Obtaining and characterising *B. napus IND* mutants

We identified two sequences from *B. napus* with >90% identity to the diploid *Brassica IND* genes (alignment in Online Resources 1). Based on the genome sequence of the *B. napus* variety Darmor-*bzh* (Chaloub et al. 2014), the accession numbers are BnaC03g32180D and BnaA03g27180D, which places them on chromosomes C3 and A3, respectively. These genes are identical to the *B. napus* orthologues identified by Braatz et al. (2018) and their annotated chromosomal locations are in agreement with our previous mapping of two *IND* genes in the Tapidor/Ningyou7 doubled haploid mapping population (Qiu et al. 2006). For simplicity, we will henceforth use the names previously assigned to these genes, *BnaC.IND.a* and *BnaA.IND.a* (Braatz et al. 2018). A phylogenetic analysis revealed that instead of being most closely related to each other, the two *B. napus IND*-like sequences cluster separately with *BraA.IND.a* and *BolC.IND.a* from the diploid progenitors (Fig. 3a). Together, these results demonstrate that the two *IND* sequences identified in *B. napus* originate from each of the two diploid progenitors (*B. rapa* and *B. oleracea*) and are not caused by duplication after the hybridisation event that formed *B. napus*.

**Fig. 3.**
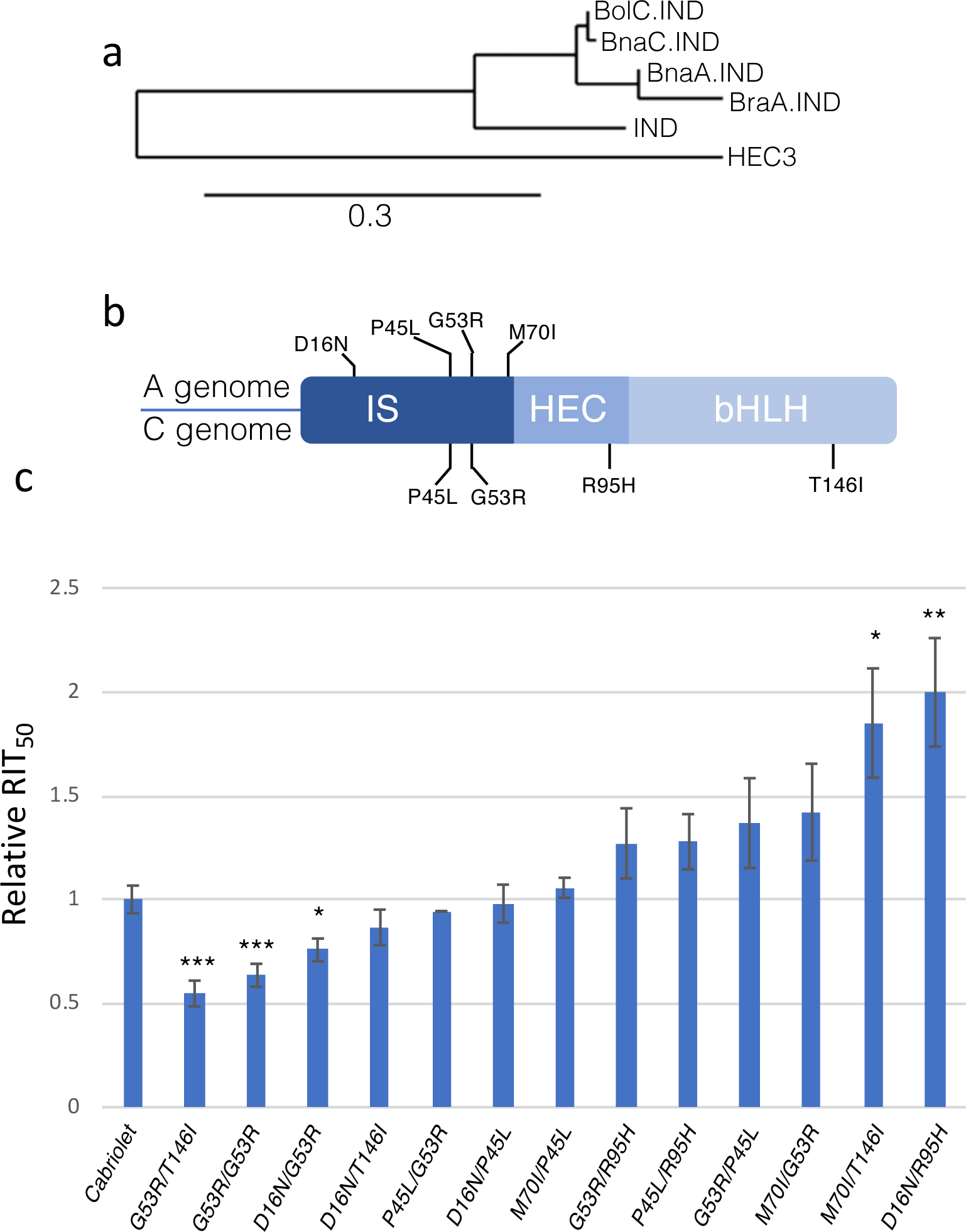
Identifying and characterising *bna.ind* mutants. **a** Phylogenetic analysis of Arabidopsis and Brassica *IND* protein sequences. HEC3 is HECATE3, which is the closest homologue of IND in *Arabidopsis* and used as an outgroup in this analysis. Scale bar indicate substitutions per site. **b** Schematic of IND protein divided into its three domains: the IND-Specific (IS), HECATE (HEC) and basic Helix-Loop-Helix (bHLH) domains. Position and nature of missense mutations are shown with four from the A genome (top) and four from the C genome (bottom). **c** RIT half lives (RIT_50_) plotted for the wild-type variety ‘Cabriolet’ and the combinations of *bna.ind* double mutants indicated in the graph with substitutions on the A genome in front of substitutions on the C genome. *, p<0.05; **, p<0.01; p<0.001.

In order to obtain loss-of-function *bnaA.ind.a* and *bnaC.ind.a* mutants, we first designed paralogue-specific amplicons for the two *B. napus IND* genes (Online Resource 2a,b) and subsequently screened the *B. napus* TILLING resource available in the variety Cabriolet from the RevGenUK platform at the John Innes Centre (www.revgenuk.ac.uk). We did not identify any mutations that would generate premature stop codons; however, out of the allelic series that were obtained, we chose four mutant alleles for each paralogue for further studies (Fig. 3b). These alleles were chosen based on the nature of the amino acid change (e.g. charge, hydrophobicity, size) and conservation among IND from *Arabidopsis* (see alignment in Online Resource 1).

In agreement with functional redundancy of the two *B. napus IND* genes, we did not observe any phenotypic defects in single mutants (Online Resource 2c,d). Crosses were therefore performed in order to obtain double mutants. We performed all combinations of crosses resulting in 16 different double-mutant lines (Online Resource 3a).

The mutant lines did not display the same dramatic effects on valve margin formation as the knock-out and over expression lines shown in Fig. 2. However, given that complete indehiscence will not be desirable for oilseed rape farmers in terms of recovering seeds, fine-tuning IND and GA4 activities through pairing of mutant alleles may provide levels of dehiscence better suited for yield increase. To quantify pod-shatter resistance of fruits from the *B. napus* mutant lines generated here, we used the Random Impact Test (RIT) assay first developed by Morgan et al. (1998). The RIT applies mechanical force to mature and dry Brassica fruits by shaking them in a container in the presence of metal ball bearings for a certain amount of time (Online Resource 4a-c). The time it takes for half of the fruits in the container to break open is taken as a measure of their level of pod shatter resistance (Morgan et al. 1998) and referred to here as the RIT_50_ value (Online Resource 4d). To better analyse this data, we developed the software package, podshatteR. Based within the free R software environment it provides a simple GUI with a step-by-step process for improved data quality analysis, curve fitting and therefore a more precise quantification of the RIT_50_ value (Online Resource file).

For the *bnaA.ind.a bnaC.ind.a* double mutants, we obtained fruits suitable for the RIT assay for 13 out of the 16 combinations and tested them in comparison to the wild-type Cabriolet variety, which is the background for the TILLING population. For simplicity, we will refer to these double mutants by their amino acid substitution as in Fig. 3c where the first annotation refers to the substitution on the A genome and the second on the C genome. The graph in Fig. 3c shows a wide range of shatter resistance from lines such as *G53R/T146I* that exhibit less resistance than wild type to *M70I/T146I* and *D16N/R95H* that are significantly more resistant than wild type. It is not clear, which residue mutations provide the effect as combinations with each of the individual mutations in the resistant lines also found in combinations with less resistance (Fig. 3c).

We analysed sections of the *D16N/R95H* and *M70I/G53R* at different developmental stages late in fruit development (Fig. 4; Online Resource 3b). Although the differences are subtle compared to the effects seen in full *ind* knockouts (Fig. 2), there are features, which may explain the increased shatter resistance in the mutant lines. At stage 16, valve margin cells are becoming apparent in wild-type fruits as small light blue cells. The definition of these cells is less pronounced in the two best performing mutant combinations (Fig. 4 left; Online Resource 3b). At stage 17B, when lignification is visible, the light-blue separation layer appears wider in the wild type (Fig. 4 middle). Moreover, at full maturity (stage 17c), valves detach completely in the wild-type fruits, but remain partially connected in the mutants (Fig. 4 right; Online Resources 3b).

**Fig. 4.**
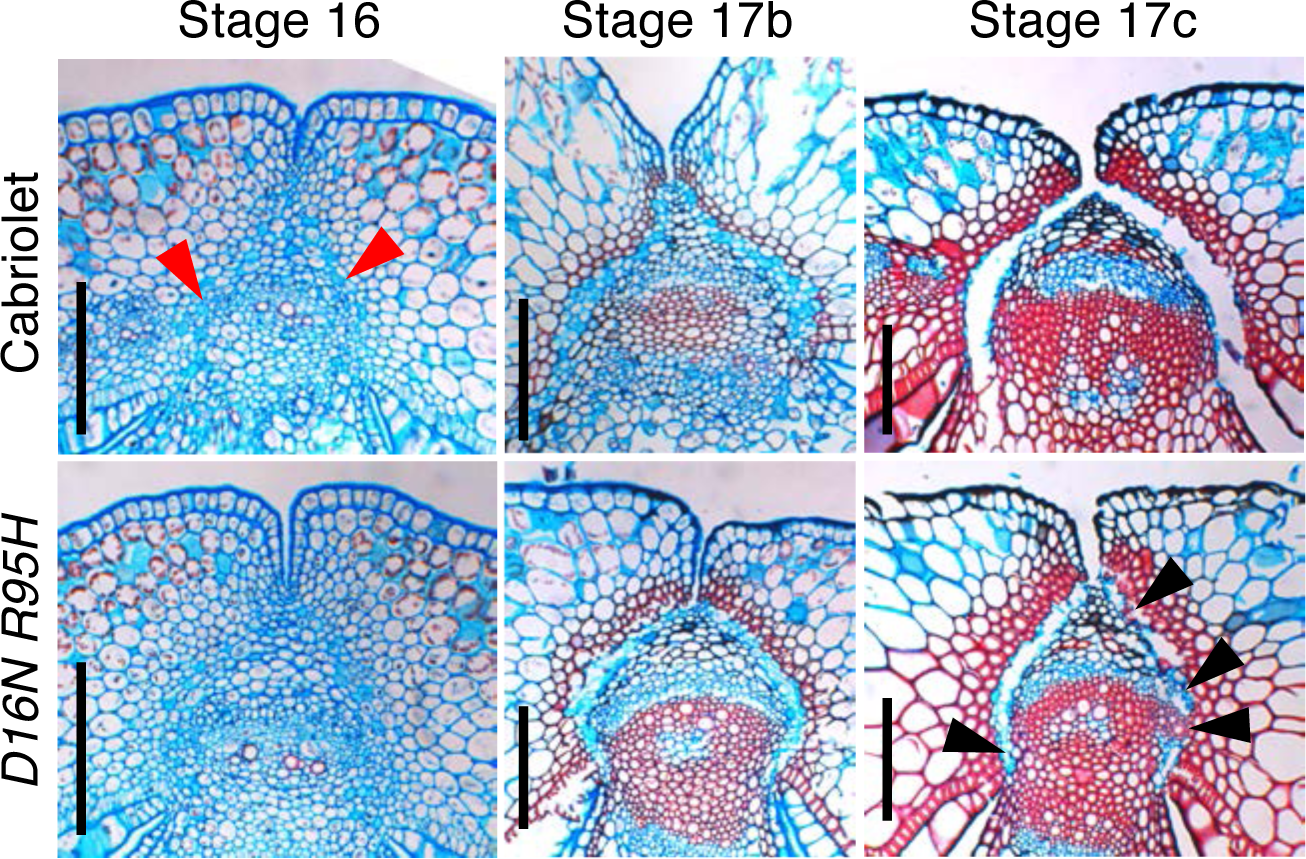
Tissue sections of fruits from Cabriolet (wild type) and the *bnaA.ind.a-D16N bnaC.ind.a-R95H* double mutant (*D16N/R95H*) at the indicated developmental stages. Red arrowheads point to separation layer primordium cells in Cabriolet. Black arrowheads indicate lack of complete separation in the *D16N/R53H* double mutant. *Scale bars* correspond to 100 µm.

Taken together, these data show that by obtaining an allelic series of mutations in the *Bna.IND* genes it is possible to make combinations in order to fine tune the level of pod shattering.

### Obtaining *B. napus GA4* mutants

We have previously described how knocking out two *GA4* paralogues in *B. oleracea* by CRISPR/Cas9 led to the production of pod-shatter resistant siliques (Lawrenson et al. 2015). Moreover, the plants displayed a dwarf architecture in agreement with the phenotype of *ga4* mutants in *Arabidopsis* (Chiang et al. 1995).

In parallel with the isolation of *B. napus ind* mutants, we attempted to obtain mutants in the *B. napus GA4* orthologues. The *B. napus* genome contains four *GA4* paralogues (BnaA06g10250D, BnaA09g57140D, BnaC05g11920D, BnaC08g38810D), which we rename for simplicity as *BnaA6.GA4, BnaA9.GA4, BnaC5.GA4* and *BnaC8.GA4*, respectively. The encoded proteins show a very high level of identity to each other and to GA4 from *Arabidopsis* of 85-90% (see alignment in Online Resource 5). Nevertheless, a phylogenetic analysis demonstrated that each of these are most closely related to the paralogues on the corresponding chromosomes in the *B. rapa* and *B. oleracea* diploids (Fig. 5a). This analysis demonstrates that *B. napus GA4* paralogues are conserved from the diploid progenitors and have not undergone any additional duplication or gene loss since the hybridisation of the diploids which formed the allotetraploid *B. napus*.

**Fig. 5.**
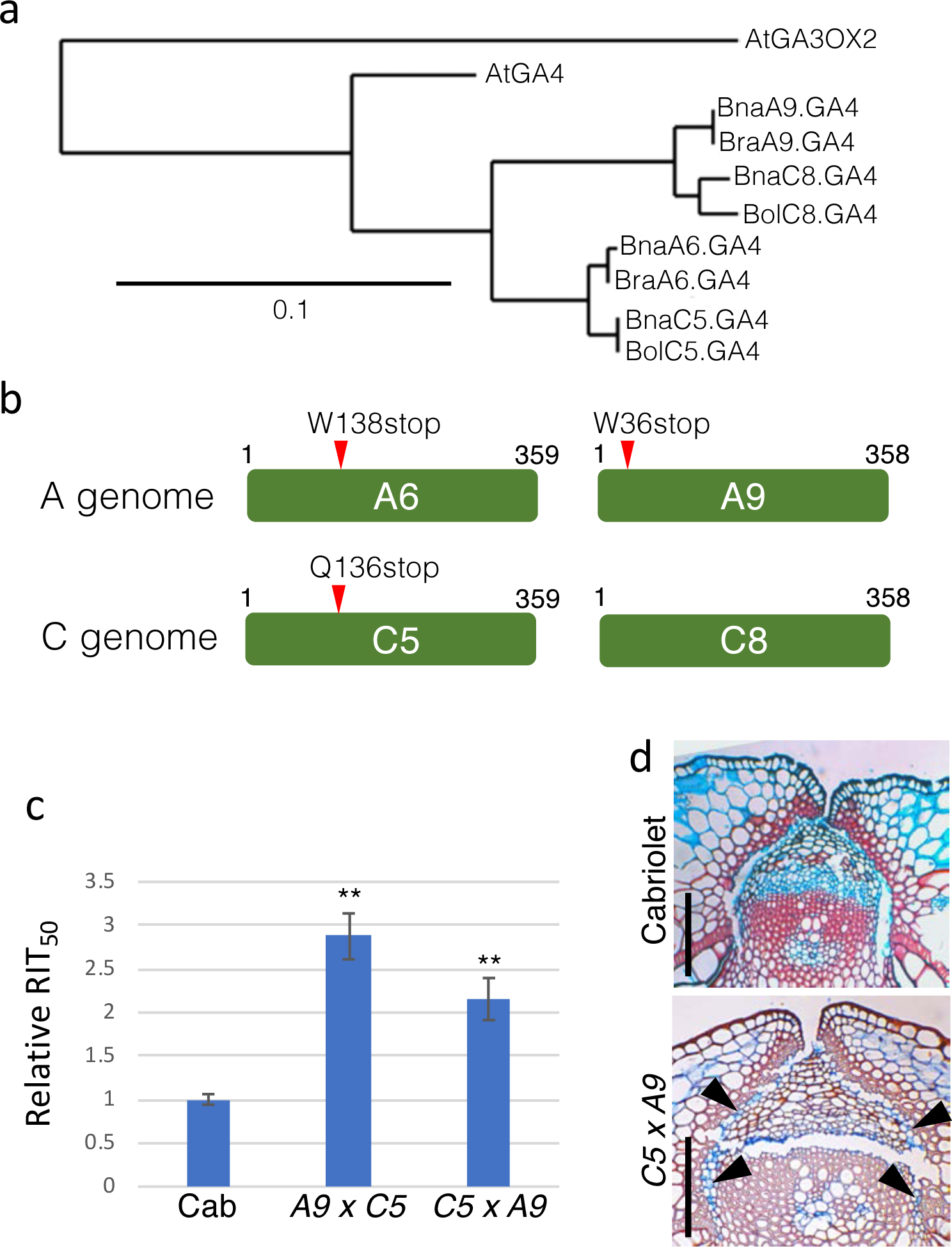
Identifying and characterising *B. napus ga4* mutants. **a** Phylogenetic analysis of Arabidopsis and Brassica *IND* protein sequences. AtGA3OX2 was used as an outgroup in this analysis. Scale bar indicates substitutions per site. **b** Schematic of the four GA4 proteins indicating position of nonsense mutations in the paralogues on chromosomes A6, A9, and C5. **c** RIT_50_ values plotted for ‘Cabriolet’ (Cab) and reciprocal crosses of the double mutant of paralogues on chromosomes A9 and C5. **, p<0.01. **d** Tissue sections of fruits from Cabriolet (wild type) and the *bnaA09.ga4 bnaC05.ga4* (*C5 x A9*) double mutant. Black arrowheads indicate lack of complete separation in the *C5 x A9* double mutant *Scale bars* correspond to 100 µm.

We designed specific amplicons for each of the four paralogues as described for the *B. napus IND* alleles above. By screening the *B. napus* Cabriolet TILLING population, we obtained allelic series of all. For three of these *GA4* genes, at least one allele gave rise to a premature stop codon in the 5’ half of the coding region (Fig. 5b, Online Resource 5) and we decided to focus on these for the subsequent analyses. As expected, due to redundancy among the paralogues, none of the single mutants led to any phenotypic defects (Online Resource 6). Hence, all three combinations of double mutants were produced by crossing and homozygous lines obtained.

The effect on valve margin development in *ga4* mutants in *Arabidopsis* is specific to the separation layer and the level of indehiscence of *ga4* fruits is therefore not as pronounced as for fruits from the *ind* mutant that have lost both separation layer and lignified layer (Arnaud et al. 2010). Rather than developing a large allelic series using combinations of missense mutations, we focussed on the nonsense mutations, which gave rise to premature stop codons in three of the *B. napus GA4* paralogues.

In a RIT analysis, we found that the combination of the *bnaA9.ga4 bnaC5.ga4* double exhibited the largest effect with shatter resistance ∼2.5 fold above wild type with significant increase in shatter resistance observed for both directions of crossing; *i.e.* when pollen from the *C5* allele was applied to *A9* stigma and vice versa (Fig. 5c). In agreement with the increased resistance, cross sections of stage-17C fruits from the *C5xA9* double mutant exhibited reduced separation at the valve margin compared to wild type (Fig. 5d).

In both examples shown here, 2-3-fold higher shatter resistance was obtained. Interestingly, this is in the same range as the best performing *bna.ind* double mutant line obtained in the ‘Express’ variety (∼3-fold increase compared to wild type) (Braatz et al. 2018). Also, a 2.5-fold improvement was obtained in the InVigor hybrid line developed by Bayer in which the two *bna.ind* mutants are heterozygous knock-outs (European Patent Specification 2008).

In conclusion, the RIT_50_ data for certain of the mutant combinations obtained here revealed a significant increase in resistance compared to wild type. In particular, the *bna.ind D16N/R95H* and *bnaA9.ga4 bnaC5.ga4* mutants led to resistance that is comparable to previous reports (Braatz et al. 2018; European Patent Specification 2008) and therefore hold promising potential for inclusion in future breeding programmes. We have recently demonstrated that higher temperature accelerates pod shatter in fruits from a range of Brassicaceae species including *Arabidopsis* and *B. napus*. Our experiments revealed this effect to be mediated at least partially by upregulation of the *IND* gene (Li et at. 2018). It is therefore plausible that the lines generated here will maintain their high shatter resistance independent of temperature increase, potentially leading to an even higher relative improvement compared to the Cabriolet wild type.

### Perspectives

In the last three decades, investment in research to understand the growth and development of *Arabidopsis thaliana* has been second to none in the plant kingdom. The amount of knowledge on *e.g.* gene regulation, hormone dynamics, metabolism and disease resistance emerging from this investment has been impressive. Nevertheless, politicians, funders and tax payers have a right to question whether this enormous attention to a small weed is justified.

Scientifically, discoveries in *Arabidopsis* have no doubt led to a significantly increased basic understanding of plant biology and more widely biology in general. Whilst this may not have immediate societal impact, it contributes to a fundamental part of being human – *i.e.* a continued strive towards understanding the world we live in.

The model-to-crop project described here provides a reassuring example that, in terms of crop improvement, the focus on *Arabidopsis* has been a valuable investment. The results in the *B. napus* crop are based purely on a pipeline originating from fundamental discoveries in *Arabidopsis* going back >20 years since the first gene was discovered (*FUL*; Gu et al., 1998). In fact, identification of the key regulators would almost certainly not have been achievable directly in a polyploid crop, such as *B. napus*. The forward screens involved and requirements to overcome redundancy of paralogues would simply make it unfeasible to identify the key regulators and elucidate the genetic and hormonal networks in *B. napus*; even with the advanced gene-editing and genomics technologies available today.

The strategy for addressing pod shatter in oilseed rape by the approach described here has depended entirely on a translational approach. This work therefore demonstrates how knowledge obtained in a model system can be successfully transferred for use in crop improvement programmes. It should therefore serve as an encouraging example for scientists, the industry and funding agencies to get involved and support such undertakings.

Vast amounts of genetics and genomics resources are becoming available directly in crop species such as wheat (Uauy, 2017), rice (Wing et al. 2018), maize (Nannas, Dawe 2015) and also in *Brassica* species (Lawrenson et al. 2015; Chalhoub et al. 2014). Therefore, addressing fundamental questions is no longer restricted to *Arabidopsis* or other relatively simple genetic systems. Scientists can now test hypotheses directly in crops to an extent that has not previously been possible, which will lead to an acceleration in crop improvement. Nevertheless, the small size, simple genetics and short generation time will ensure that *Arabidopsis thaliana* and other model plants remain attractive and important model systems for future discoveries with the potential to save billions from starvation and which no doubt will expand our fundamental understanding of biology on the planet.

## Author contributions

PS, NS and LØ designed the research. NP developed podshatteR software. PS, NS, MB, MI, CO and RW performed experiments. L.Ø. wrote the manuscript and all authors commented on it.

## Acknowledgements

We thank the following science support services at the John Innes Centre: RevGenUK TILLING platform, Bio-imaging, Photography and Horticulture for skilful assistance. This work was supported by grants BB/J533055/1 (Follow-on Fund), BB/I017232/1 (Crop Improvement Research Club) and BB/P003095/1 (Strategic LoLa) to L.Ø. from the Biotechnological and Biological Sciences Research Council and by the Institute Strategic Programme grants BB/J004553/1 and BB/P013511/1 to the John Innes Centre.

## Online Resources

### Online Resource 1

Multiple alignment of IND protein sequences from Arabidopsis and Brassica species including HEC3 from Arabidopsis as an outgroup. Alignment was done using ClustalOmega. Mutated residues used in the analysis of the manuscript are indicated by a black circle.

### Online Resource 2

**a** Schematic outline of *IND* gene showing position of paralogue-specific primers (BnaC-f and BnaA-f) and BnaAC-r, which will amplify both. **b** Agarose gel showing paralogue-specific PCR amplification using primers BnaC-f/BnaAC-r to amplify *BnaC.IND.a* (IND-C) or BnaA-f/BnaAC-r to amplify *BnaA.IND.a* (IND-A) on gDNA from *B. rapa* (A), *B. oleracea* (C) and *B. napus* (A/C). **c** Effect of single mutations in *bna.ind* genes. SEM images of mature (stage 17) fruits from Cabriolet (wild type) and indicated *bna.ind* single mutants. **d** Tissue sections of mature (stage 17c) fruits from Cabriolet (wild type) and indicated *bna.ind* single mutants. *Scale bars* correspond to 1 mm in c and 100 µm in d.

### Online Resource 3

**a** Table of double-mutant combinations. ‘X’ indicates fruits were obtained for RIT while ‘-’ indicates no RIT was performed on those lines due to lack of fruits. **b** Tissue sections of fruits from Cabriolet (wild type) and the *bnaA.ind.a-M70I bnaC.ind.a-G53R* (*M70I/G53R*) double mutant at the indicated developmental stages. Red arrowheads point to separation layer primordium cells in Cabriolet. Black arrowheads indicate lack of complete separation in the *M70I/G53R* double mutant. *Scale bars* correspond to 100 µm.

### Online Resource 4

Random Impact Test analysis and visualisation of the podshatteR software output. **a** RIT machine on lab bench. **b** photo of sample with ball bearings prior to shaking. **c** photo of sample after one round of 8-sec shaking. **d** example of two samples tested in triplicates with number of remaining intact pods plotted against time of shaking. Values are fitted with decay curves calculated by software generated here. RIT_50_ values are calculated as the time it takes to break 10 pods out of 20.

### Online Resource 5

Multiple alignment of GA4 protein sequences from Arabidopsis and Brassica species. Alignment was done using ClustalOmega. Mutated residues used in the analysis of the manuscript are indicated by a black circle.

### Online Resource 6

Effect of mutations in *bna.ga4* genes. SEM (upper row) and tissue section (lower row) images of mature (stage 17) fruits from Cabriolet (wild type), *bnaA.ga4* and *bnaC.ga4* single mutants. Red arrowhead indicates valve that is opening, black arrowheads point to separation layer in cross sections. *Scale bars* correspond to 1 mm for SEMs and 100 µm for sections.

### Online Resource 7

Oligonucleotides used in this study.

