## Supplemental Information for "The power of model-to-crop translation illustrated by reducing seed loss from pod shatter in oilseed rape"

### Online Resource 1

```
HEC            -----MNNYNMNPSLFQNYTWNNIINSSNNNNK--NDDHHHQHNNDPIGMAMDQYTQLHI
IND            MMEPQPHHLLMD--------WNKANDLLTQEHAAFLNDPHHLMLDPP-------------
BraA.IND.a     MME--HHHLLMN--------WNKPIDLITEENS--FNHNPHFIVDPP-------------
BnaA.IND.a     MME--HHHLLMN--------WNKPIDLITEENS--FNHNPHFIVDPP-------------
BolC.IND.a     MME--PHHLLMN--------WNKPIDLITQENS--FNHNPHFMVDPP-------------
BnaC.IND.a     MME--PHHLLMN--------WNKPIDLITQENS--FNHNPHFMVDPP-------------
                     ::  *:        **:  :  .:::    :   *   : *

HEC            FNPFSSSHFPPLSSSLTTTTLLSGDQEDDEDEEEPLEELGAMKEMMYKIAAMQSVDIDPA
IND            --PETLIHLD--------------------EDEEYDEDMDAMKEMQYMIAVMQPVDIDPA
BraA.IND.a     --SETLSHFQPPPTIFSDHG--GGEEAEEEEEEEGEEEMDPMKKMQYAIAAMQPVDLDPA
BnaA.IND.a     --SETLSHFQPPPTIFSDHG--GGEEAEEEEEEEGEEEMDPMKKMQYAIAAMQPVDLDPA
BolC.IND.a     --SETLSHFQPPPTVFSDHG--GGEEA---EDEEGEEEMDEMKEMQYAIAAMQPVDIDPA
BnaC.IND.a     --SETLSHFQPPPTVFSDPG--GGEEA---EDEEGEEEIDEMKEMQYAIAAMQPVDIDPA
                 . :  *:                     ::**  *::. **:* * **.**.**:***

HEC            TVKKPKRRNVRISDDPQSVAARHRRERISERIRILQRLVPGGTKMDTASMLDEAIRYVKF
IND            TVPKPNRRNVRISDDPQTVVARRRRERISEKIRILKRIVPGGAKMDTASMLDEAIRYTKF
BraA.IND.a     TVPKPNRRNVRVSDDPQTVVARRRRERISEKIRILKRMVPGGAKMDTASMLDEAIRYTKF
BnaA.IND.a     TVPKPNRRNVRVSDDPQTVVARRRRERISEKIRILKRMVPGGAKMDTASMLDEAIRYTKF
BolC.IND.a     TVPKPNRRNVRVSEDPQTVVARRRRERISEKIRILKRMVPGGAKMDTASMLDEAIRYTKF
BnaC.IND.a     TVPKPNRRNVRVSEDPQTVVARRRRERISEKIRILKRMVPGGAKMDTASMLDEAIRYTKF
               ** **:*****:*:***:*.**.*******.****:*:****:**************.**

HEC            LKRQIR-LLNNNTGYTPPPPQDQASQAVTTSWVSPPPPPSFGRGGRGVGELI
IND            LKRQVR-ILQPHSQIGAPMANPSYL-----CYYHNSQP--------------
BraA.IND.a     LKRQVR--------LASSASHSAWS-----SYV-------------------
BnaA.IND.a     LKRQVRLLLQPHTQLGAPMSDPSCL-----CYYHNSDT--------------
BolC.IND.a     LKRQVR-LLQPHTQLGAPMSDPSRL-----CYYHNSDT--------------
BnaC.IND.a     LKRQVR-LLQPHTQLGAPMSDPSRL-----CYYHNSDT--------------
               ****:*          .. .          .:
```

**Online Resource 1.** Multiple alignment of IND protein sequences from Arabidopsis and Brassica species including HEC3 from Arabidopsis as an outgroup. Alignment was done using ClustalOmega. Mutated residues used in the analysis of the manuscript are indicated by a black circle.

### Online Resource 2

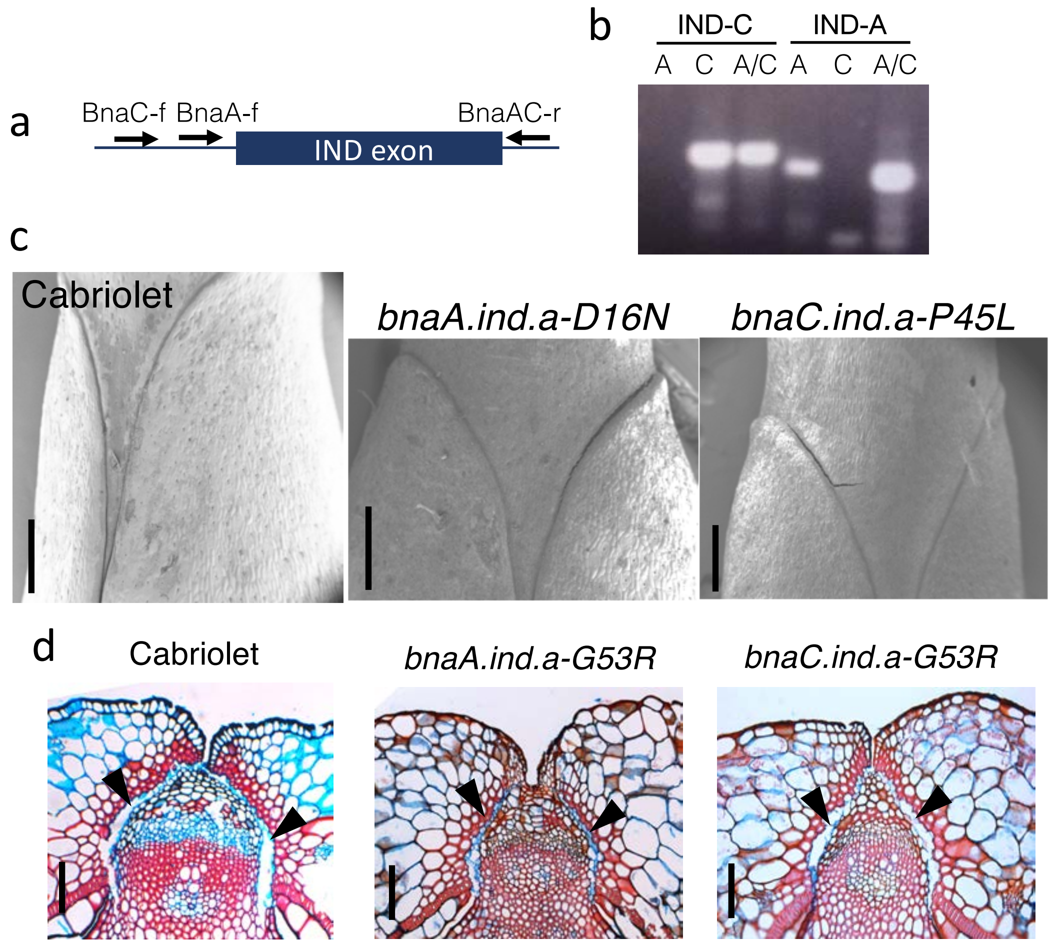

**Online Resource 2. a** Schematic outline of *IND* gene showing position of paralogue-specific primers (BnaC-f and BnaA-f) and BnaAC-r, which will amplify both. **b** Agarose gel showing paralogue-specific PCR amplification using primers BnaC-f/BnaAC-r to amplify *BnaC.IND.a* (IND-C) or BnaA-f/BnaAC-r to amplify *BnaA.IND.a* (IND-A) on gDNA from *B. rapa* (A), *B. oleracea* (C) and *B. napus* (A/C). **c** Effect of single mutations in *bna.ind* genes. SEM images of mature (stage 17) fruits from Cabriolet (wild type) and indicated *bna.ind* single mutants. **d** Tissue sections of mature (stage 17c) fruits from Cabriolet (wild type) and indicated *bna.ind* single mutants. *Scale bars* correspond to 1 mm in c and 100 µm in d.

Online Resource 3

a

| *bnaA.ind.a* \ *bnaC.ind.a* | P45L | G53R | R95H | T146I |
|---|---|---|---|---|
| D16N | X | X | X | X |
| P45L | - | X | X | - |
| G53R | X | X | X | X |
| M70I | X | X | X | X |

b

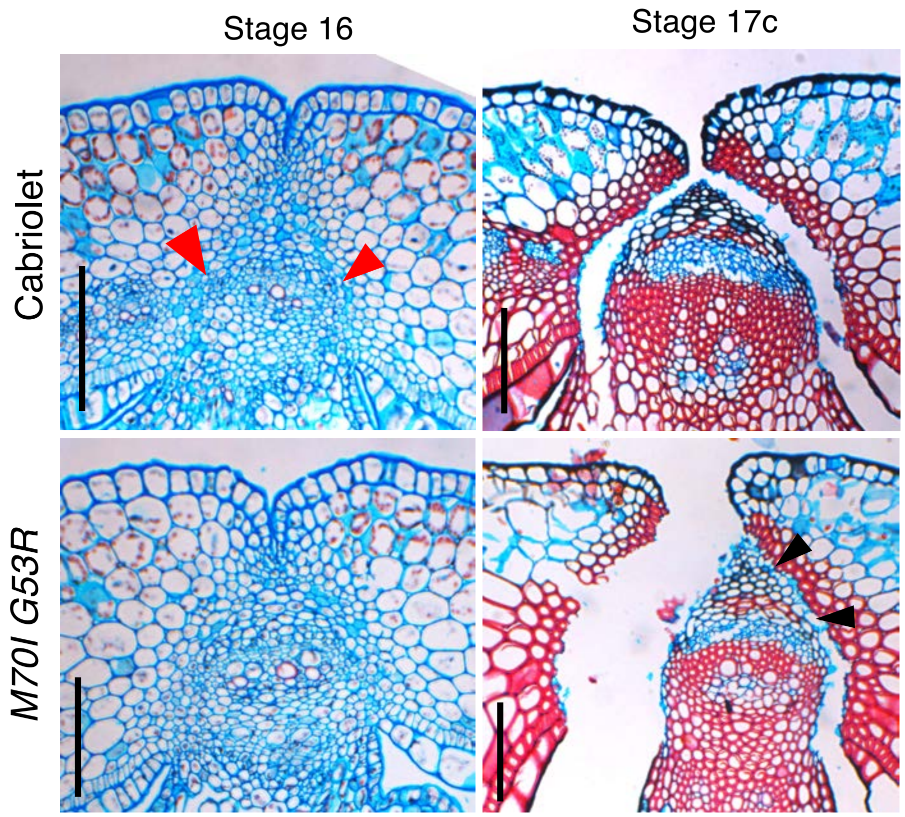

**Online Resource 3. a** Table of double-mutant combinations. 'X' indicates fruits were obtained for RIT while '-' indicates no RIT was performed on those lines due to lack of fruits. **b** Tissue sections of fruits from Cabriolet (wild type) and the *bnaA.ind.a-M70I bnaC.ind.a-G53R* (*M70I/G53R*) double mutant at the indicated developmental stages. Red arrowheads point to separation layer primordium cells in Cabriolet. Black arrowheads indicate lack of complete separation in the *M70I/G53R* double mutant. *Scale bars* correspond to 100 µm.

Online Resource 4

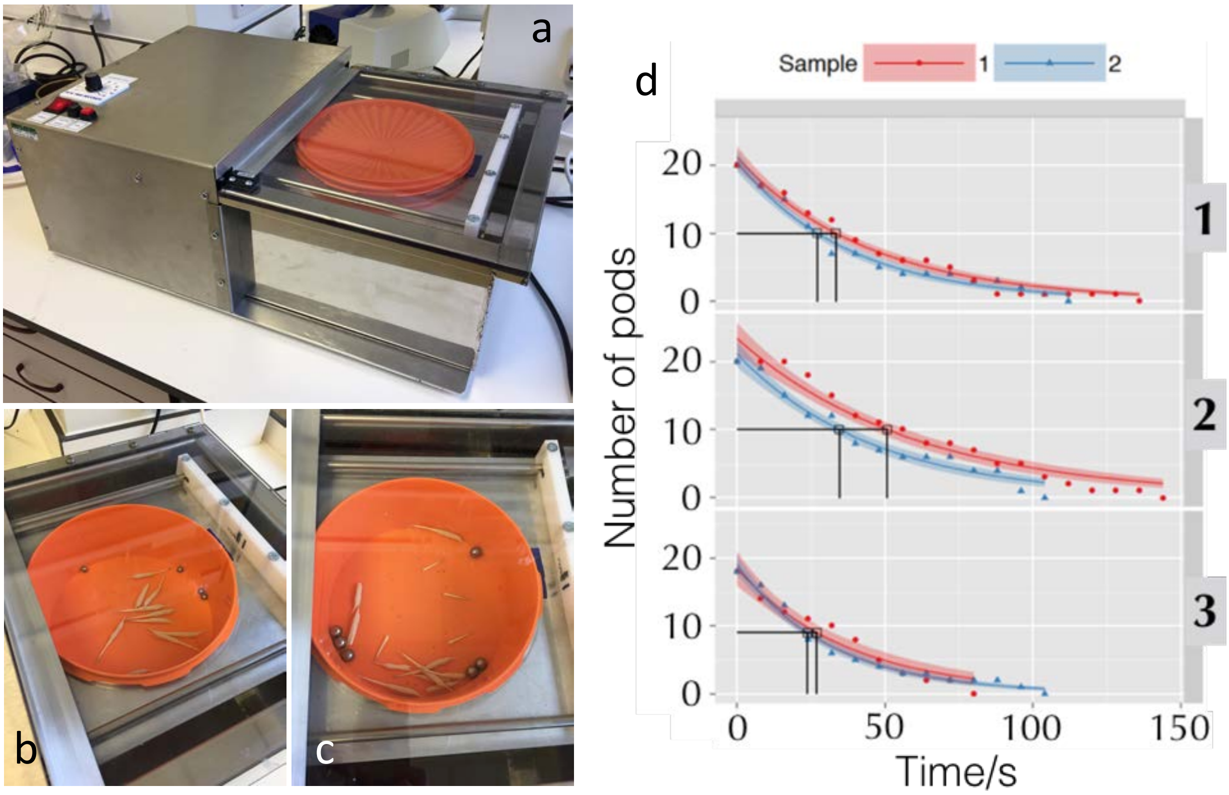

**Online Resource 4.** Random Impact Test analysis and visualisation of the podshatteR software output. **a** RIT machine on lab bench. **b** photo of sample with ball bearings prior to shaking. **c** photo of sample after one round of 8-sec shaking. **d** example of two samples tested in triplicates with number of remaining intact pods plotted against time of shaking. Values are fitted with decay curves calculated by software generated here. $RIT_{50}$ values are calculated as the time it takes to break 10 pods out of 20.

### Online Resource 5

```
AtGA4           MPAMLTDVFRGHPIHLPHSHIPDFTSLRELPDSYKWTPKDDLLFSAAPSPPATGENIPLI
BnaC5.GA4       MPTMLTDVFRGHPIHLPHSHQPDFTSLSELPDSYTWTPKDDPLLDAAPSPPAASENIPLI
BnaA6.GA4       MPTMLTDVFRGHPIHLPHSHQPDFTSLSELPDSYTWTPKDDPLLDAAPSPPAASENIPLI
BnaC8.GA4       MPTVLTDVFRGHPIHLPHTHQPDFTSLSELPDSYTWTSKDDPLFTAPPSPPGAGESIPLI
BnaA9.GA4       MPTVLTDVFRGHPIHLPHTHQPDFTSLSELPDSYTWTSKDDPLFTAPPSPPDAGESIPLI
                **::**************:* ****** ******.** *** *: * **** :.*.****

AtGA4           DLDHPDATNQIGHACRTWGAFQISNHGVPLGLLQDIEFLTGSLFGLPVQRKLKSARSETG
BnaC5.GA4       DLNHPDAANQIGSACRTWGAFQIANHGVPLELLQGIEFLTGSLFQLPVHRKLKAARSETG
BnaA6.GA4       DLNHPDAANQIGSACRTWGAFQIANHGVPLELLQGIEFLTGSLFQLPVQRKLKAARSETG
BnaC8.GA4       DLNHPDAANQIGRACRTWGAFQIANHGVPLELLQDIEFLTGSLFQLPVQHKLEAARSDAG
BnaA9.GA4       DLNHPDAANQIGRACRTWGAFQIANHGVPLELLQGIEFLTGSLFQLPVQRKLKAARSDTG
                **:****:**** **********:****** ***.********* ***::**::***::*

AtGA4           VSGYGVARIASFFNKQMWSEGFTITGSPLNDFRKLWPQHHL-NYCDIVEEYEEHMKKLAS
BnaC5.GA4       FSGYGVARISSFFNKQMWSEGFTITGSPLNDFRKLWPQHHLNNYCDIVEEYEEQMQKLAS
BnaA6.GA4       FSGYGVARISSFFNKQMWSEGFTITGSPLNDFRKLWPQHHLNNYCDIVEEYEEQMQKLAS
BnaC8.GA4       FSGYGVARISSFFNKKMWSEGFTITGSPLNDFRKLWPQLHL-NYCDIVEQYEEQMQKLAS
BnaA9.GA4       FSGYGVARISSFFNKKMWSEGFTITGSPLNDFRKLWPQLHL-NYCDIVEQYEEQMQKLAS
                .********:*****:********************** ** *******:***:*:****

AtGA4           KLMWLALNSLGVSEEDIEWASLSSDLNWAQAALQLNHYPVCPEPDRAMGLAAHTDSTLLT
BnaC5.GA4       KLMWLSLTSLGVSEEDIKWASANSDSDWAQSALQLNHYPVCPEPDRAMGLAAHTDSTLLT
BnaA6.GA4       KLMWLSLTSLGVSEEDIKWASANPDLNWAQSALQLNHYPVCPEPDRAMGLAAHTDSTLLT
BnaC8.GA4       KLMWLSLNSLGVSEEDIKWARVSSDLNWAQSALQLNHYPVCPEPDRAMGLAPHTDSTLLT
BnaA9.GA4       KLMWLSLNSLGVTEEDIKWATVSSDLNWAQSALQLNHYPVCPEPDRAMGLAPHTDSTLLT
                *****:*.****:****:**  . * :***:******************** ********

AtGA4           ILYQNNTAGLQVFRDDLGWVTVPPFPGSLVVNVGDLFHILSNGLFKSVLHRARVNQTRAR
BnaC5.GA4       ILHQNNTAGLQVFRDDLGWVTVPPVPGSLVVNVGDLFHILSNGLFKSVLHRARVNQTRSR
BnaA6.GA4       ILHQNNTAGLQVFRDDLGWVTVPPVPDSLVVNVGDLFHILSNGLFKSVLHRARVNQTRSR
BnaC8.GA4       ILYQNNTAGLQVFRDDLGWVTVPPVPGSLVVNVGDLFHILSNGLFKSVIHRVRVNQTRPR
BnaA9.GA4       ILYQNNTAGLQVFRDDFGWVTVPPVPGSLVVNVGDLFHILSNGLFKSVIHRARVNQTRPR
                **:*************:*******.*.*********************:**.****** *

AtGA4           LSVAFLWGPQSDIKISPVPKLVSPVESPLYQSVTWKEYLRTKATHFNKALSMIRNHREE
BnaC5.GA4       LSVAFLWGPQSDIKISPVPKLVSPVGSPLYRSVTWTEYLRTKATHFNEALSMIRNHIDE
BnaA6.GA4       LSVAFLWGPQSDIKISPVPKLVSPVGSPLYRSVTWTEYLRTKATHFNEALSMIKNDIDE
BnaC8.GA4       LSVAFLWGPRSDTKISPVPKLVSPDESPLYRSVTWTEYLRTKATHFNKALSMIRNHREK
BnaA9.GA4       LSVAFLWGPRSDTKISPVPKLVSPDESPLYRSVTWTEYLRTKATHFNKALSMIRNHREK
                *********:** ***********  ****:****.***********:*****:*. ::
```

**Online Resource 5.** Multiple alignment of GA4 protein sequences from Arabidopsis and Brassica species. Alignment was done using ClustalOmega. Mutated residues used in the analysis of the manuscript are indicated by a black circle.

Online Resource 6

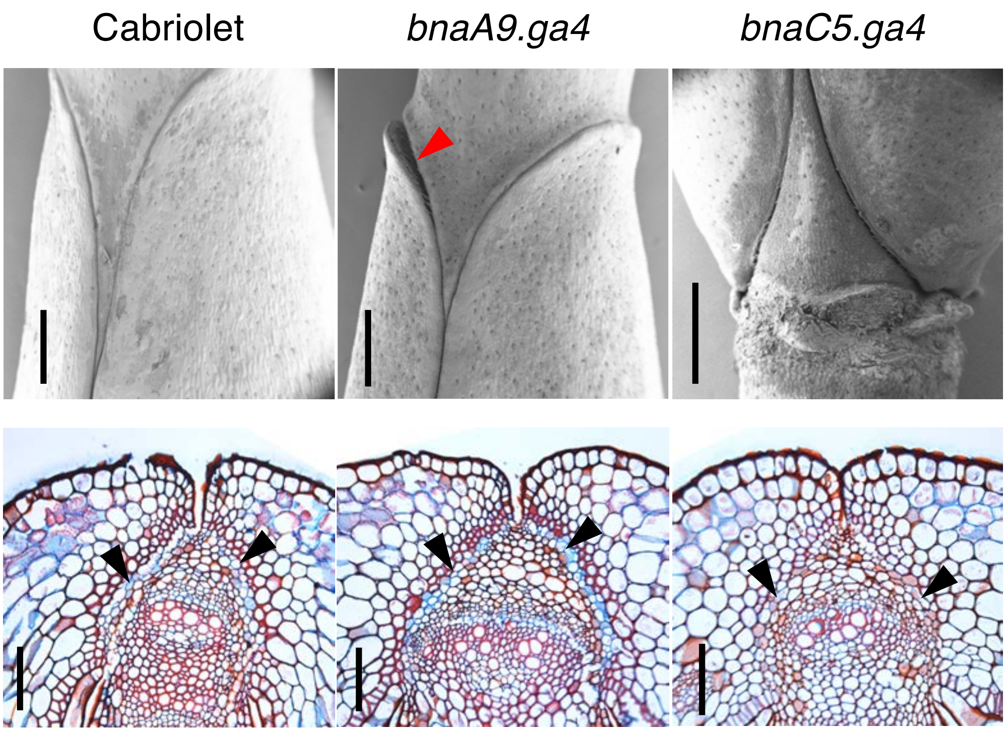

**Online Resource 6.** Effect of mutations in *bna.ga4* genes. SEM (upper row) and tissue section (lower row) images of mature (stage 17) fruits from Cabriolet (wild type), *bnaA.ga4* and *bnaC.ga4* single mutants. Red arrowhead indicates valve that is opening, black arrowheads point to separation layer in cross sections. *Scale bars* correspond to 1 mm for SEMs and 100 µm for sections.

Online Resource 7

Oligonucleotides used in this study

***Bna.GA4***

| | |
|---|---|
| A6-F | gtccaacacttccataatcttcc |
| A6-R | GAATGACATGGCGAAATCTCTGT |
| C5-F | gactctccgactcttccataatt |
| C5-R | ACATGGCAAATTTTCGGTTTCT |
| C8-F | gacacaaaacatctatcgaat |
| C8-R | gccgtctttagcatatgtaaattgagccg |
| A9-F | ccaaaacaagagcttaaacatatgc |
| A9-R | gcaaccaatatatatcatgacc |

***Bna.IND***

| | |
|---|---|
| IND A-F | aacattcatacacgcactac |
| IND A-R | atcaacatgaaacgcgtgat |
| IND C-F | ccgcaaatacaaacatatttagt |
| IND C-R | atcaacatgaaacgcgtgat |

**Online Resources**

**The podshatteR software**

podshatteR is a browser tool or web app for plotting and analysing pod shatter data collected from Random Impact Tests (RIT) assays (see Materials and Methods). This app works for data that may have repeated samples and/or used a randomised block design.

It facilitates visualisation of measured pod shatter RIT data and determine the 'half-life' ($RIT_{50}$) of each sample using a curve fitting routine in R.

podshatteR is available at https://github.com/nstjhp/podshatteR

**Generation of podshatterR software**

podshatteR is released as a collection of R scripts and interacted within a browser through the shiny package. Launch the web app with the command

```
R -e "shiny::runApp('~/shinyAppFolderLocation')"
```

After launching the web browser and navigating to the correct port you will see the application landing page which explains the details of how to use the tool. On the first tab the tool can read in your input RIT data formatted as described on the homepage in a csv file. A plot of the data is produced to allow visual inspection of any errors or ambiguities that can then be resolved before proceeding. The following tab calculates the fit to the data. The model used is an exponential decay function to keep the number of free parameters minimal i.e.

$$Value = Aexp(-b \times Time)$$

where A is the intercept (which should be similar to the starting number of pods), and b is the decay rate. Through propagation of errors confidence intervals are computed for each sample. The next tab allows for the plotting of the half-life locations and the ability to download a csv file containing all the calculated half-lives.

The half-life is calculated for each sample as the value at which half the initial number of pods has shattered, following the model fit. In the final tab you can download a subset of the plots in PDF format. The x-axis can be fixed across the rows/columns, or can be fixed, and the final plot size of the downloaded PDF can be chosen by the user.

The podshatteR code was built with R version 3.1.2 and uses the following packages (versions):

- shiny (0.11.1)
- ggplot2 (1.0.0)
- grid (3.1.2)
- reshape2 (1.4.1)
- plyr 1.8.1
- minpack.lm (1.1-8)
- propagate (1.0-4)

**Using the podshatteR software**

In order to use podshatteR, a desktop PC equipped with Statistical package R (version 3.2 or later) is required. The app is written in R using the `shiny` package for display in the browser and was built using the following versions:

- `R` v3.1.2
- `shiny` v0.11.1
- `ggplot2` v1.0.0
- `grid` v3.1.2
- `reshape2` v1.4.1
- `plyr` v1.8.1

- `minpack.lm` v1.1-8
- `propagate` v1.0-4

**Step-by-step procedure**

1. After the installation of R, the following packages will be required to run the App. Type (or copy from the INSTALL file within the podshatteR folder) the command below, to install the required packages:

install.packages(c("shiny", "ggplot2", "grid", "reshape2", "plyr", "minpack.lm", "propagate"), dependencies=TRUE)

2. If running for the first time, restart R, then type the command to load the Shiny R package:

>library(shiny)

3. In the File dropdown menu, change the working directory to the podshatteR folder. To open and run the application in a web browser, type the command:

>runApp()

The web app is split up into a number of sections, which can be navigated using the bar at the top.

4. Select the Upload and plot data tab

- Click `Browse...` to find and upload data. This will have to be saved as a .CSV file in the required format (shown on the About tab). Please note capital letters are required at the start of each column header, failure to provide this will prevent data from being loaded.
- This tab will then read the data, display them in a table, and plot the data with lines to facilitate spotting of errors in the data entry.
- The plots show every line with repeated samples in different colours and symbols.

If using more than one block, then the separate blocks will be shown in separate sub-tabs.

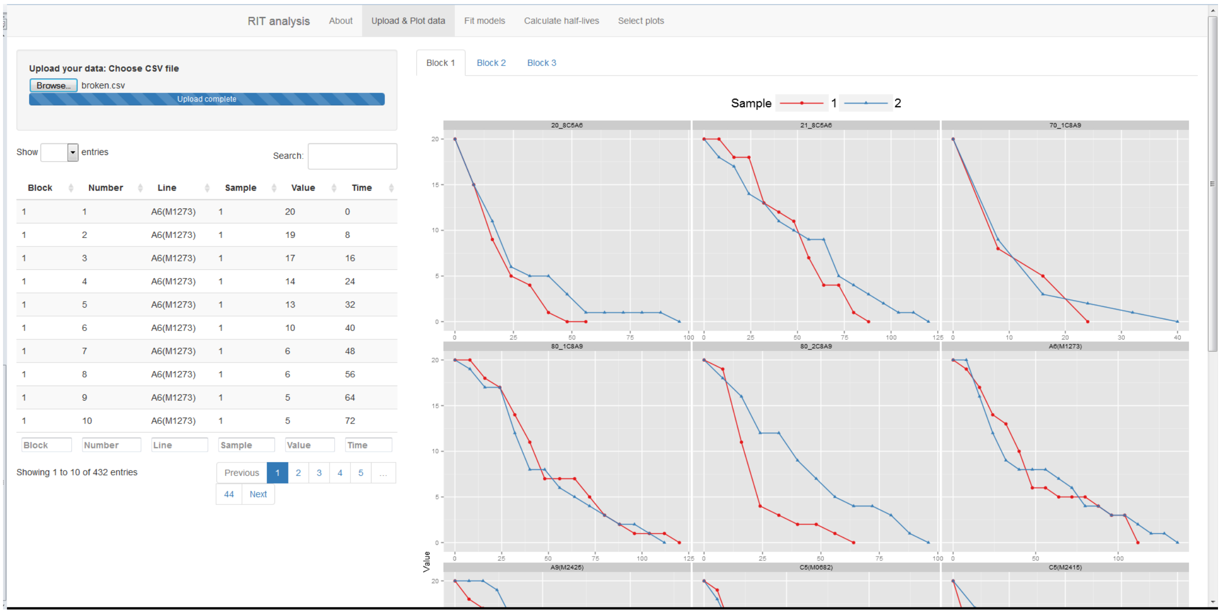

Fig.1 An example of data plotted in the upload and plot data tab.

5. Select the Fit models tab and press the Fit models button. For every combination of block, line and sample the model must be fitted and confidence intervals calculated. A plot will be displayed showing the raw data points and a line for the fitted model with a semi-transparent ribbon showing the confidence intervals.

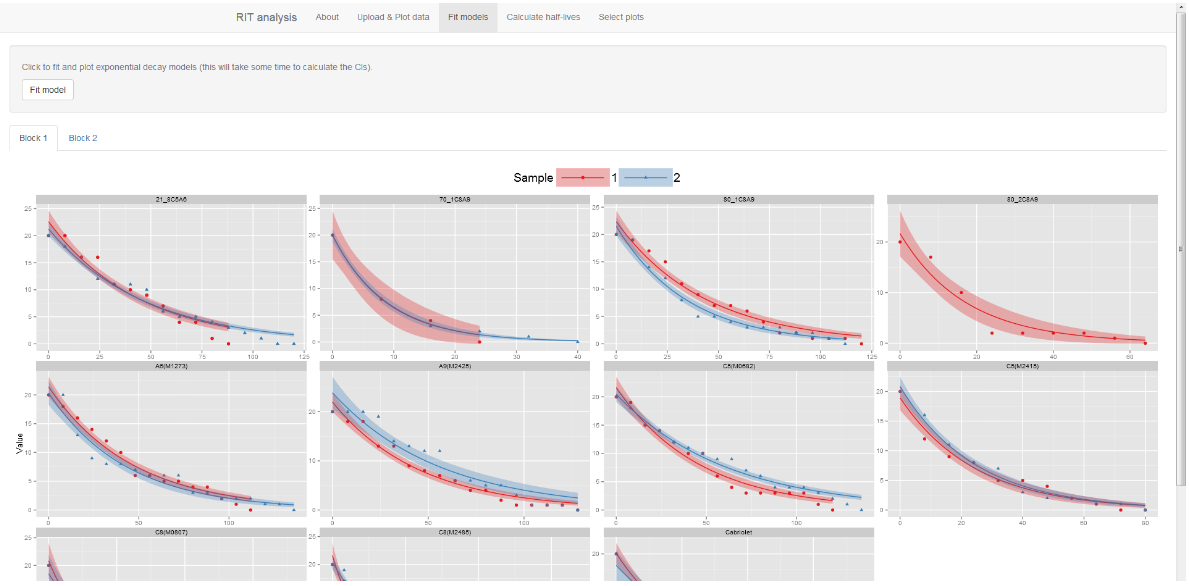

Fig 2. An example of the Fit models tab showing plotted curves and confidence intervals

6. Select the Calculate half-lives and click the button. A table will appear giving the result for each combination of variables at which half the initial pods are estimated to have shattered, along with half the number of initial pods. Plots will also appear building on the previous model fit plots with dark lines showing where this half-life point is. A .CSV file of these data can be downloaded using the download button.

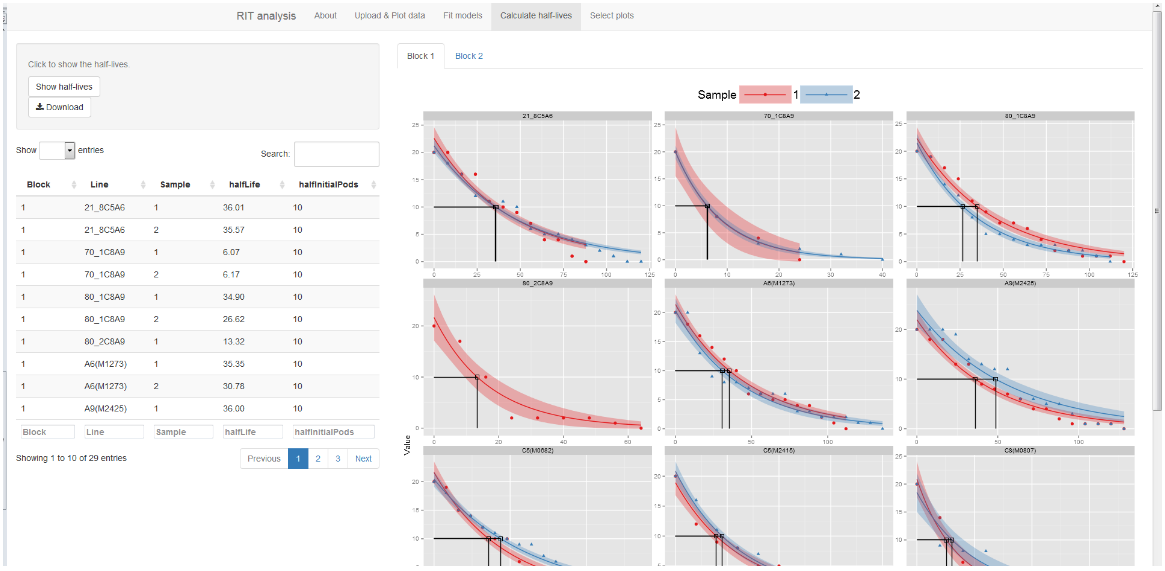

Fig. 3 The output from the Calculate half-lives tab showing half-life estimate table (left) and half-life plotted on the fitted curves (right).

7. Select plots tab allows simple selection of a subset of plots to display for presentations etc. by checking the boxes and clicking `Show plots`. If new plots are selected, simply check different boxes and click `Show plots` again.

Each panel can be chosen to have the same axes or each one to be different. With the same axes the panels will all be scaled to the most-resistant line so you can easily select the ones that shatter at noticeably different times for instance.

To allow full control of plot size, adjust the `Plot height` and `Plot width` sliders. These are reactive so there is no need to click on `Show plots` again (numbers represent inches for the downloaded figure).

Clicking `Download` will facilitate downloading of a PDF of the current figure (some aspects, particularly the legend size, may differ between screen display and saved plot, but this can be adjusted using the sliders).
